# How weedy *Arabidopsis thaliana* dominated the world: ancestral variation and polygenic adaptation

**DOI:** 10.1101/2024.04.28.591542

**Authors:** Cheng-Yu Lo, Chih-Cheng Chien, Ya-Ping Lin, Pei-Min Yeh, Cheng-Ruei Lee

## Abstract

The composition of a species could change with demographic turnovers, where dominant populations quickly expanded and replaced others. However, whether such events have a genetic basis remains to be investigated. Previous studies showed that *Arabidopsis thaliana* experienced a significant demographic turnover, where “non-relicts” replaced “relicts” throughout Eurasia. Here, we showed that non-relicts have smaller seeds, more seeds per fruit, and a higher germination rate, making them more competitive over relicts. Using a unique population enriching relict alleles while minimizing population structure, we identified candidate loci and showed that such trait divergence was caused by the divergent sorting of multiple ancient haplotypes in a Mendelian gene and joint allele frequency change of polygenes affecting single-trait divergence and multi-trait covariance. This study is one of the few genetic investigations of species-wide demographic turnover, emphasizing the importance of processes different from the much-focused hard selective sweep.

## Introduction

Population structure exists in most species, and the composition of a species is often affected by the dynamics among these populations. One of the most dramatic forces might be large demographic turnover, in which a strongly competitive population quickly expanded and replaced other native populations. One of the best examples would be the replacement of Neanderthals by anatomically modern humans [1, 2]. In this case, however, the genetic investigation of the underlying causes remains difficult.

In addition to humans, the model plant *Arabidopsis thaliana* is one of the few cases of these large-scale demographic turnovers. Recent studies have identified several highly diverged populations inhabiting southern Europe, Africa, and southwestern China (the relicts), and other worldwide accessions are descendants of a population originating from Central Europe (the non-relicts) [3-6]. While the relicts were once diverse and widespread throughout Eurasia, the non-relicts expanded worldwide, quickly replacing and hybridizing with local relicts [5, 6]. It was hypothesized that the expansion of this population might be associated with the spread of agriculture, analogous to human commensal weeds [5, 7]. It remains unclear, however, what phenotypic and genetic characteristics were responsible for the initial success of non-relicts in replacing relicts.

Weedy plants are generally believed to have higher dispersibility, smaller stature, abundant seed production, and higher growth rates [8, 9]. Interestingly, a previous study found that seeds of Cvi (a relict) are much larger than Ler (a non-relict), while the latter has higher fecundity [10] and different hypocotyl growth in response to light [11]. We aim to investigate whether these and several other traits from previous studies represent a systematic difference between relicts and non-relicts. Using a unique natural hybridizing population between these relicts and non-relicts, we performed genome-wide association studies to identify loci associated with between-group divergence while minimizing the confounding effects from population structure. Finally, the genetic architecture and selection signatures of candidate loci were investigated.

This study systematically investigated the phenotypic, genetic, genomic, and selection architecture of within-species large-scale population turnover. We found that the relict vs. non-relict differences evolved through the divergent sorting of ancestral variations in a single gene or the allele frequency changes of many loci with minor effects on single or multiple traits. Distinct from the common emphasis on the hard selective sweep of large-effect genes on largely qualitative traits, we provided a different picture of the evolution of quantitative traits, which eventually resulted in significant demographic changes within a species.

## Results

### Phenotypic differences between relicts and non-relicts

Previous studies revealed that the relict accession Cvi has many traits that are different from the non-relict accession Ler, such as seed size, seed number, and hypocotyl length [10, 11]. To verify whether these are general differences between relicts and non-relicts, we chose 30 Eurasian non-relicts with little introgression from relicts and all 21 previously identified relicts [3, 5] for comparison in the common garden experiment (Fig. 1a; Supplementary Data 1).

**Figure 1.**
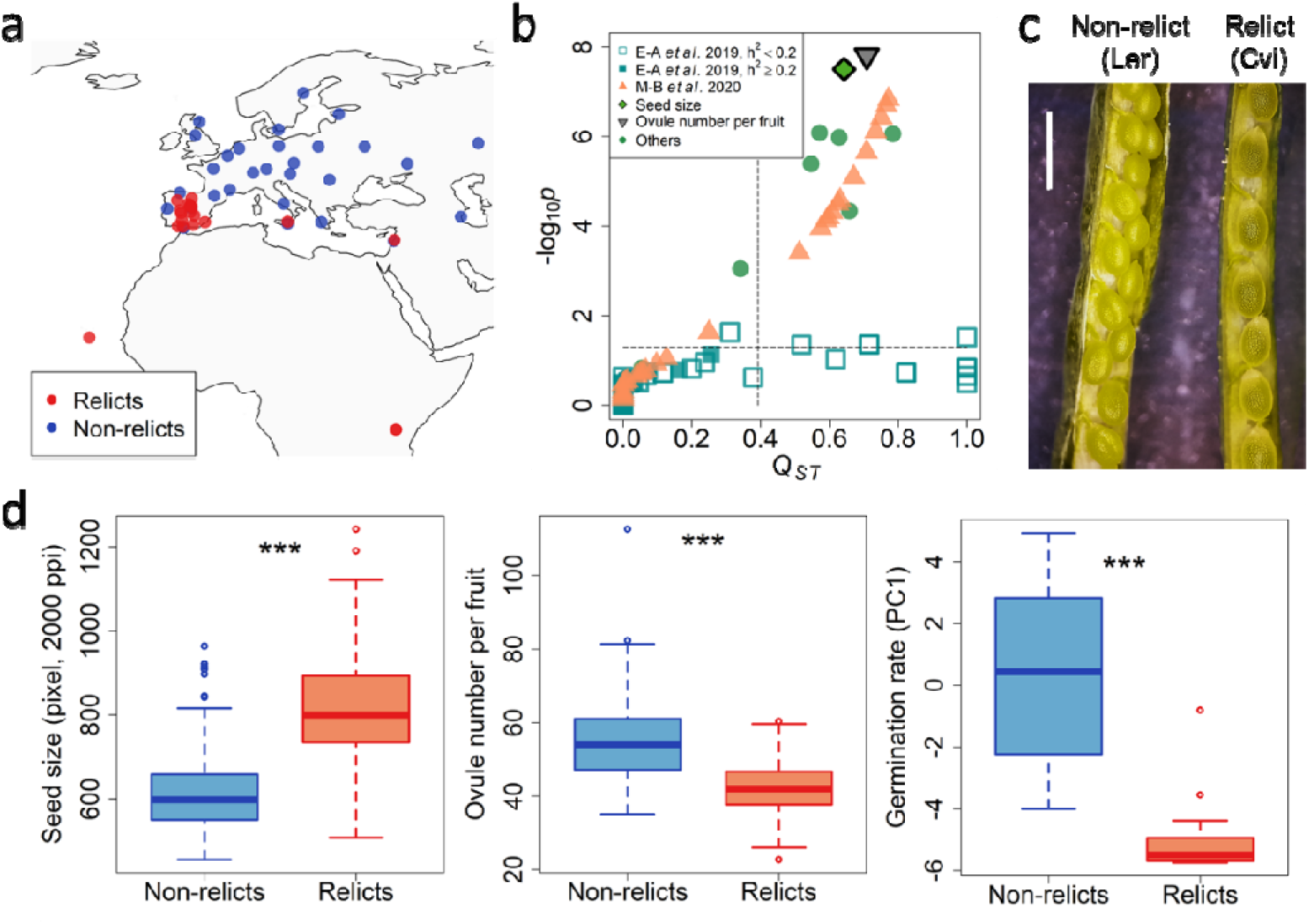
Trait divergence between relicts and non-relicts. **(a)** Accessions sampled for phenotype comparison. **(b)** Magnitudes of trait divergence in terms of *Q*_*ST*_ and *p* values in analysis of variance. Shown are the traits measured in this and two previous studies [12, 13]. For Exposito-Alonso et al. [12], traits were separated based on heritability. The vertical dashed line denotes upper 5% tail of SNP *F*_*ST*_, and the horizontal dashed line denotes *p* value of 0.05. **(c)** Seed size and the number of seeds in one representative siliques of non-relicts (left) and relicts (right). Scale bar is 1 mm. **(d)** Trait value distribution of the three target traits in this study. *** *p* < 0.001

We found significant differences in many seed- and yield-related traits (Fig. 1b; Supplementary Data 2). The seeds of relicts were larger than those of non-relicts in length and width (Fig. 1c, d). The average size of the relict seeds was approximately 820 pixels (under 2000 dpi), 33% larger than that of the non-relict seeds (615 pixels; Fig. 1d). In contrast, non-relicts have about 55 ovules in one silique, while relicts can only produce about 43 ovules. Since there was no significant difference in silique length, non-relicts appear to pack more and smaller seeds in a limited space than relicts (Fig. 1c; Supplementary Data 2). As the overall fruit number and proportion of undeveloped ovules did not differ significantly, the higher ovule number per fruit made non-relicts have a higher overall seed yield than relicts. Such trait difference was likely under strong divergent selection, as seed size and ovule number have much higher *Q*_*ST*_ values than the upper 5% tail of SNP *F*_*ST*_ distribution (0.39249; Fig. 1b; Supplementary Data 2). Other correlated traits, such as seed number per fruit and seed length and width, also have high *Q*_*ST*_. While it was known that the Cvi accession has a much longer hypocotyl than Ler [11], our results showed this was an accession-specific property rather than a systematic difference between relicts and non-relicts (Supplementary Data 2).

To widen the search for potential traits under divergent selection, wevre-analyzed data from previous studies [12, 13]. Exposito-Alonso *et al*. [12] collected survival and reproduction traits under different environments, yet there were few traits with significant differences between relicts and non-relicts. While some traits have very high *Q*_*ST*_, they tend to have very low heritability, reflecting the high micro-environmental variances under field trials and resulting in the lack of ANOVA significance (Fig. 1b; Supplementary Data 2). Martínez-Berdeja *et al*. [13] measured the germination rate of 559 accessions at 4 °C and 10 °C. The main difference between relicts and non-relicts was in the overall germination rate, mainly summarized by the PC1 (principal component 1) axis (Fig. 1b; Supplementary Data 2). The germination rate of relicts is much lower than non-relicts under all conditions (Fig. 1d), and most of the traits associated with primary dormancy (PC1) have high *Q*_*ST*_ values (Fig. 1b; Supplementary Data 2). The PC2 axis reflects the difference in germination rate between the two temperatures, yet no significant difference exists between relicts and non-relicts. These results suggested that selection favoring higher fecundity, smaller seeds, and reduced seed dormancy may have facilitated rapid non-relict expansion. Therefore, we further investigated the genetic architecture of seed size (SS), ovule number per fruit (ONF), and the PC1 of germination traits (overall germination rate).

### The genetic architecture of trait divergence

While genome-wide association studies (GWAS) were performed for seed size [14] and ovule number [15] in previous studies using a cohort of few relicts, we focused on the Iberian population (215 accessions). Being a long-term hybrid zone between relicts and non-relicts [5], Iberia possesses the genetic component from both groups, alleviating the population structure problem. Indeed, for SS, we identified a novel GWAS peak on chromosome 2 (Fig. 2a), which was not found in the previous study [14] and exhibited the difference between relicts and non-relicts (see below). However, GWAS results for ONF and germination do not have apparent peaks (Fig. 2a).

**Figure 2.**
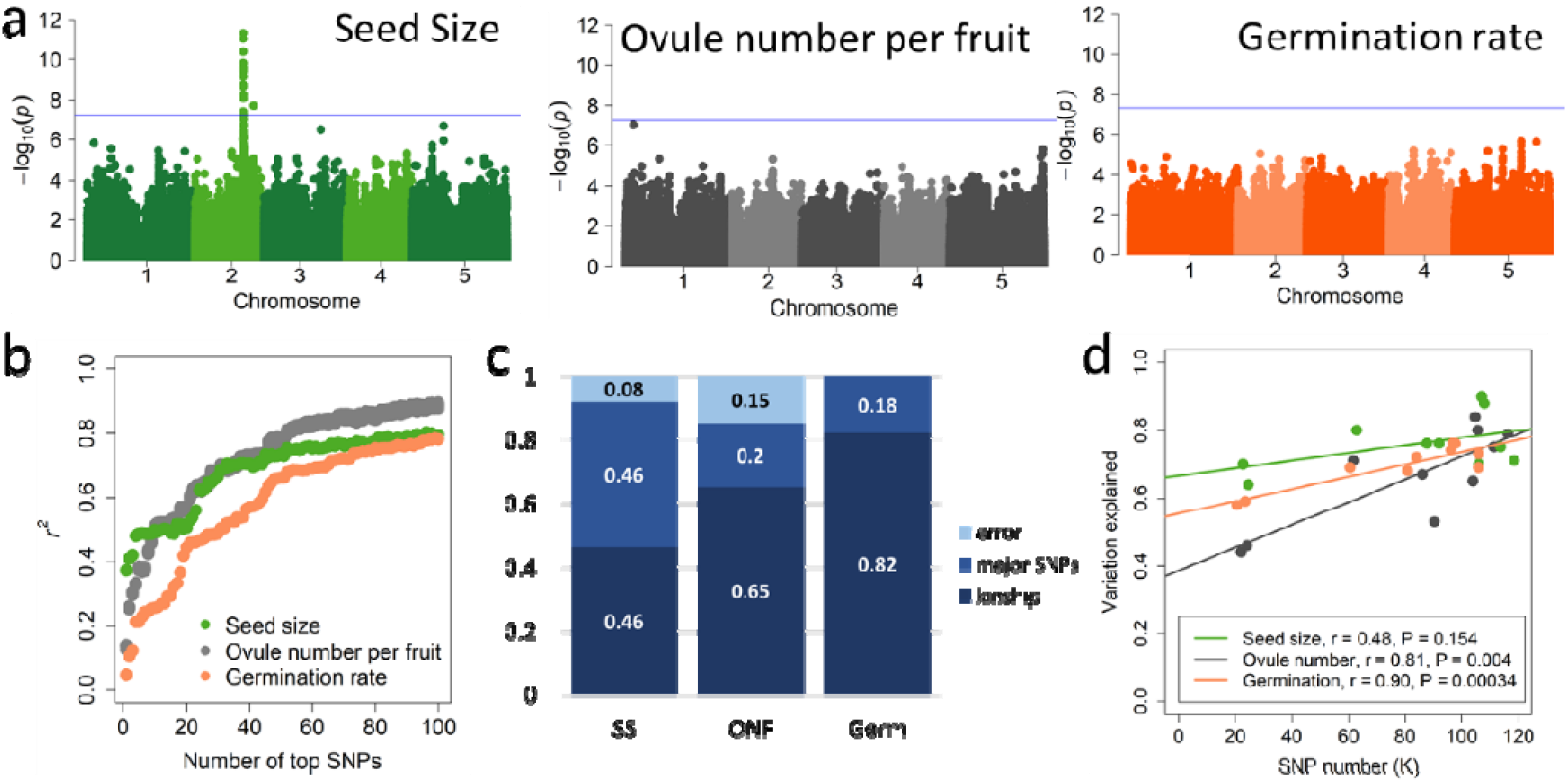
The genetic architecture of highly diverged traits between relicts and non-relicts. **(a)** Genome-wide association study (GWAS) results. The bonferroni threshold is shown by a solid line. **(b)** Ridge regression estimating proportional trait variation explained by 100 LD-pruned SNPs with highest GWAS scores. The estimation of each SNP is repeated 100 times. **(c)** Proportional trait variation explained by major SNPs (Mendelian factor) or kinship (polygenic factor), estimated by the Bayesian Sparse Linear Mixed Model (BSLMM). **(d)** Regression between trait variation explained by each chromosome arm and the number of SNPs (size) on the arm.

High kinship-heritability and *Q*_*ST*_ were observed for both ONF (*h*^*2*^ = 0.977, *Q*_*ST*_ = 0.7088) and germination (*h*^*2*^ = 0.945, *Q*_*ST*_ = 0.708), suggesting their lack of GWAS peak may be caused by the polygenic architecture. Using ridge regression on the top 100 LD-pruned SNPs with the highest GWAS scores, we showed that the most significant SNP explained almost 40% of variation for SS but much less for ONF and germination (Fig. 2b). Nevertheless, the cumulative effect of these SNPs explains about 80% variation of all traits. Further, using the Bayesian Sparse Linear Mixed Model (BSLMM) in GEMMA [16], we decomposed the proportional variation explained by major SNPs or by kinship (polygenic effects). The major SNPs explained 46% of SS, while the kinship effects had much higher contribution in ONF and germination (Fig. 2c). Finally, following the logic in many human studies [17, 18], we separated the A. thaliana genome into ten chromosome arms and estimated the trait variation explained by the cumulative effects of all SNPs in each arm (Fig. 2d). For a highly polygenic trait, there should be a strong positive correlation between chromosome arm size and variation explained. As expected, while this relationship is highly significant for ONF (*p* = 0.004) and germination (*p* < 0.001), no such pattern was observed for seed size (*p* = 0.154, Fig. 2d). In summary, SS appears to be more Mendelian, and ONF and germination are highly polygenic, explaining their lack of strong GWAS peaks.

A recent study showed accessions from northern Europe had larger seed sizes than typical non-relicts [19], potentially caused by the introgressed relict genomic regions [5]. Given the lack of the seed-enlarging allele of a major gene in the north (see below), the pattern might result from the polygenic component of relict genomes. To test this, we performed genomic prediction for seed size and ovule number per fruit of all accessions. Indeed, accessions from northern Europe have larger seeds and fewer ovules per fruit, and the predicted trait values are highly correlated with the number of relict-derived haplotypes in each accession (Supplementary Fig. 1a). The predicted trait values were validated in accessions from other studies [15, 20], exhibiting high predicted-observed correlation (Supplementary Fig. 1b).

### AT2G32590, the candidate gene affecting seed size

The seed size difference between relicts and non-relicts is mainly affected by the locus on chromosome 2 (Fig. 2a). We compared the seed size between Col-0 and mutant lines of seven genes under the GWAS peak. All mutant lines are homozygous except for the lines of AT2G32590 (*EMBRYO DEFECTIVE 2795; EMB2795*), which appears to be homozygous mutant lethal. Among all lines tested, only the two *EMB2795* heterozygous mutant lines produce much larger seeds than the wild type (Supplementary Fig. 2).

A non-synonymous coding SNP (Chr2: 13830140) in *EMB2795* is highly associated with seed size. Accessions with the reference allele (T) have smaller seed sizes than those with the alternative allele (A) (Fig. 3a). Around half of relicts and only 1% of non-relicts carry the seed-enlarging allele. This allele is concentrated in the regions where relicts occur naturally, including the Iberian Peninsula, Morocco, eastern Africa, and Yunnan, China (Fig. 3a). No consistent expression difference was observed between relict and non-relict accessions (Supplementary Fig. 3).

**Figure 3.**
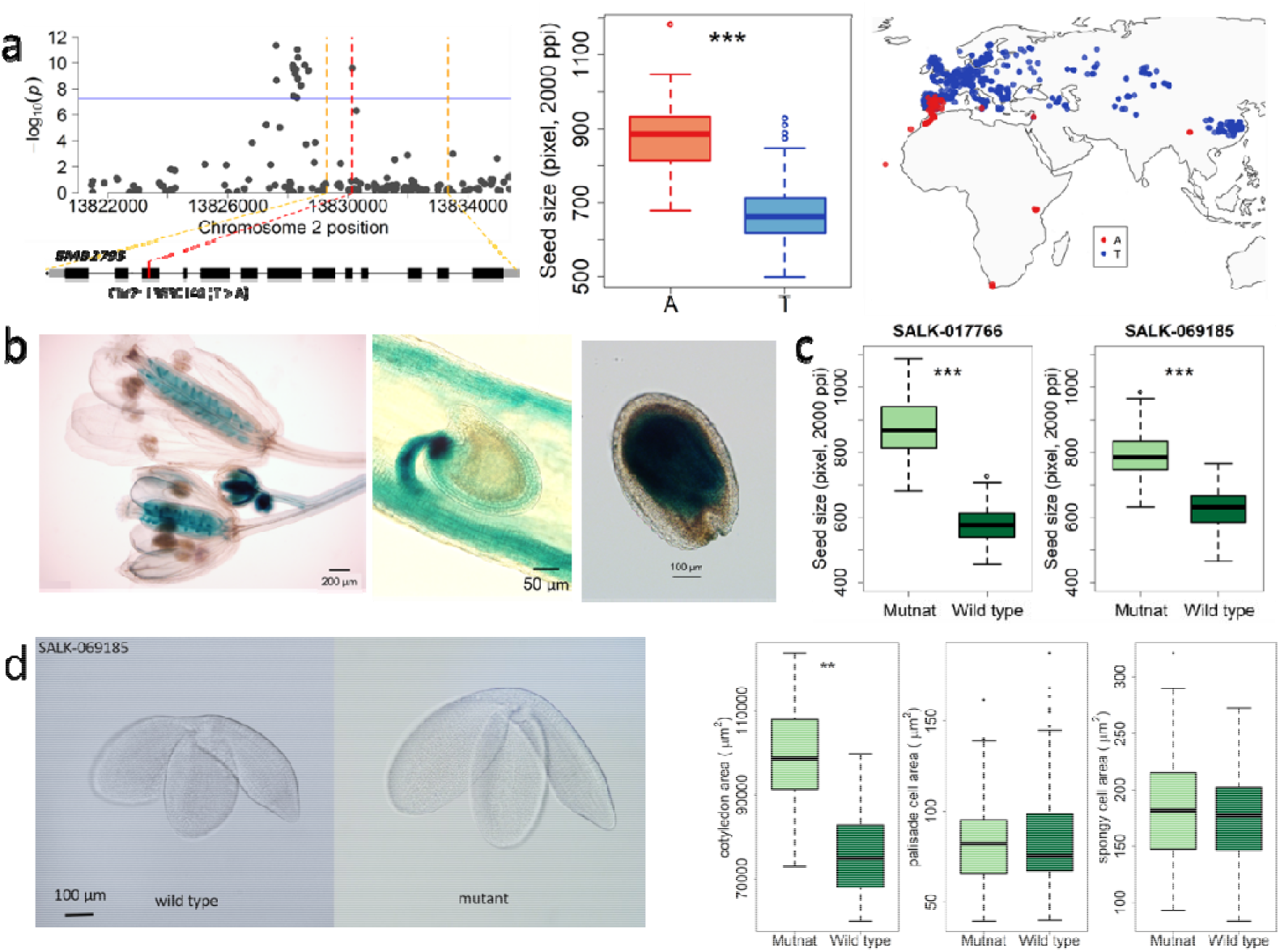
AT2G32590 (*EMB2795*) as a candidate affecting seed size differences between relicts and non-relicts. **(a)** Zoom in of genome-wide association study (GWAS) results, seeds sizes of accessions with different alleles, and the geographic distribution of the two alleles. The bonferroni threshold is shown by a solid line. **(b)** GUS staining results of *EMB2795* promoter activities in different developmental stages. **(c)** The seed size comparison between wild type and two AT2G32590 heterozygous mutant lines. **(d)** Embryo size differences between the wild type and mutant lines of AT2G32590. Shown are the photos and the size comparisons of cotyledon, palisade cell, and spongy cell. ** *p* < 0.01

*EMB2795* encodes non-SMC condensing complex subunit H. *EMB2795* is required for seed growth and development, and mutagenesis of this gene results in embryo lethality and severe defects in seed development at the pre-globular stage [21]. Our GUS reporter results showed that it was highly expressed in the buds and developing pistils but not stamens or petals of flowers. The GUS signal was also evident in the young embryo and seed coat during the stage when the seed coat enlarged to determine the final seed size. It was also expressed in the embryo during the seed-filling stage (Fig. 3b). The expression pattern was in line with the stage of rapid growth in fruit length and seed size [10, 22], and this gene expresses during the stages when the cells are actively dividing (Supplementary Fig. 4). The heterozygous mutants with lower *EMB2795* expression (Supplementary Fig. 4) tend to produce larger seeds (Fig. 3c), and their enlarged seeds also contain larger embryos (Fig. 3d). No significant difference in palisade or spongy cell size exists (Fig. 3d), suggesting the embryo size difference was caused by higher cell number. Consistently, in litchi, large-seeded accession exhibits a lower expression of the *EMB2795* homolog [23].

**Figure 4.**
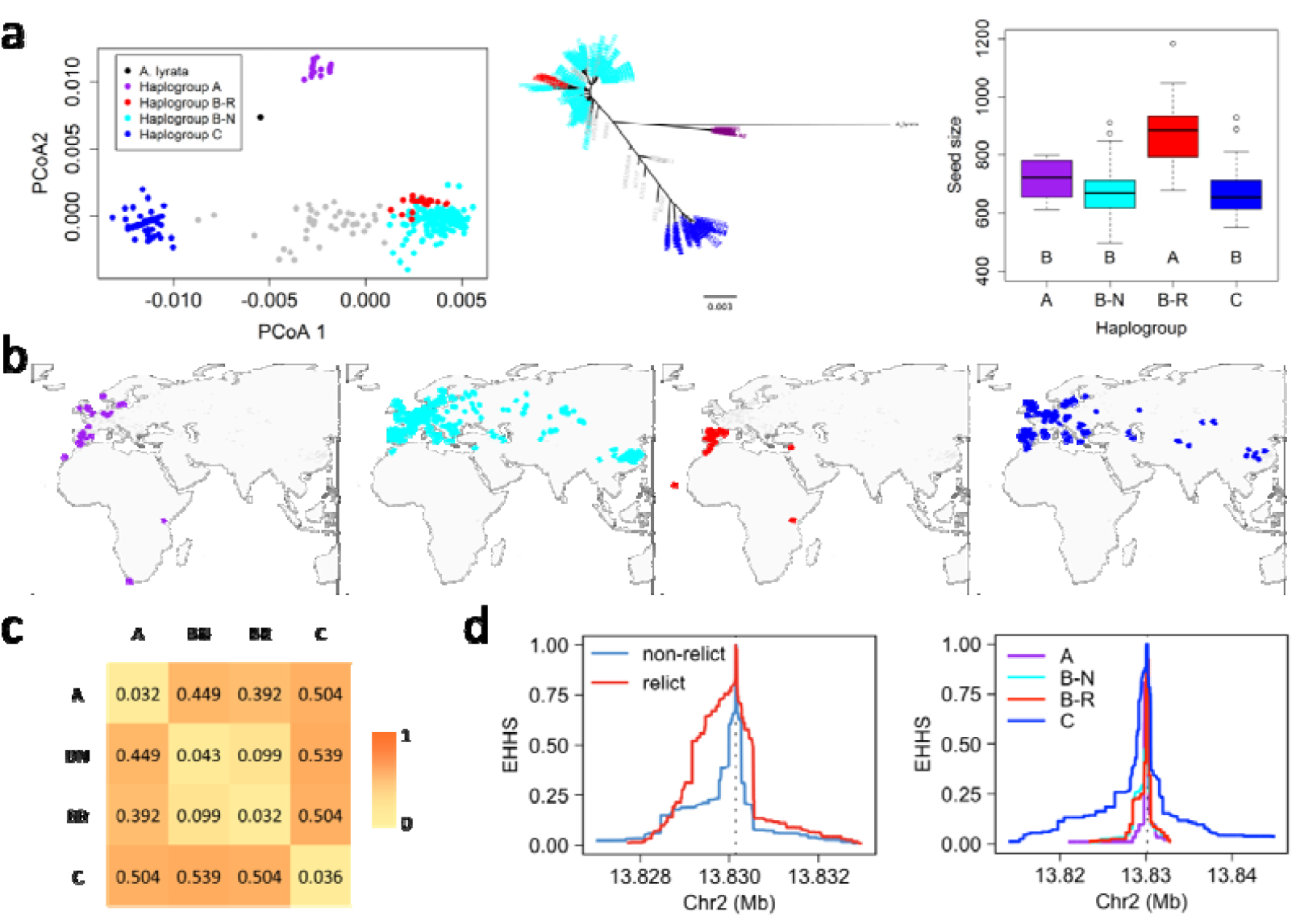
Lack of hard selective sweep for *EMB2795*. **(a)** Principal coordinate analyses (PCoA), phylogenetic tree, and seed sizes conferred by haplogroups of *EMB2795* coding region. **(b)** Geographic distribution of the four haplogroups. Colors correspond to the haplogroups in panel a. **(c)** Pairwise mean genetic differences across haplogroups (Dxy), scaled by the mean *thaliana-lyrata* divergence in this gene. **(d)** Extended haplotype homozygosity analyses comparing relicts versus non-relicts or among the haplogroups.

### The lack of signals of hard selective sweep in *EMB2795*

The *EMB2795* coding region could be separated into three highly diverged haplogroups, A, B, and C (Fig. 4a). Incorporating the diagnostic SNP (Fig. 3a), we further separated haplogroup B into B-R (for relict) and B-N (for non-relict). Only group B-R exhibited significantly larger seeds than others (Fig. 4a). In addition to B-R, haplogroup A was likely associated with relicts, as it was distributed in Africa, southwestern Europe, and northwestern Europe, consistent with previous inferences that northern accessions also contain genomic segments from relicts (Fig. 4b). Although not statistically significant, accessions with haplogroup A also had slightly larger seeds. On the other hand, haplogroups B-N and C were found across Eurasia, consistent with the pan-Eurasia range expansion of non-relicts [6]. While large variation exists among accessions with the two non-relict haplogroups (B-N and C), their seed sizes are highly correlated with the number of relict-derived haplotypes in the genome (Supplementary Fig. 5), consistent with the polygenic contribution from relict genetic components in addition to this large-effect locus.

**Figure 5.**
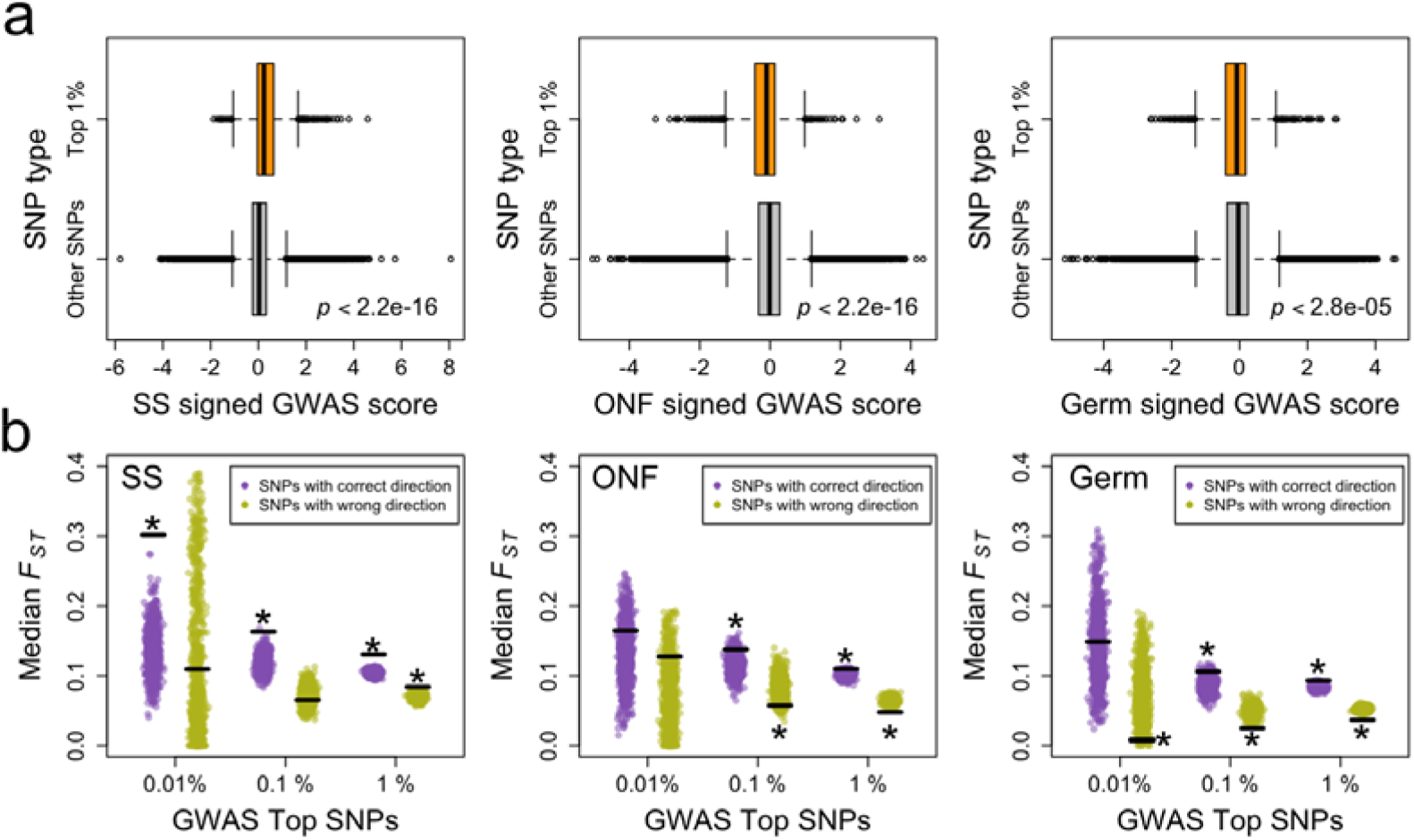
Signatures of polygenic adaptation on single traits. **(a)** Comparison of GWAS scores for SNPs with the top 1% *F*_*ST*_ value versus other SNPs. The GWAS scores were polarized so that SNPs whose relict allele decreases trait values have negative values. **(b)** Comparison of *F*_*ST*_ values for SNPs with the top 0.01%, 0.1%, or 1% GWAS scores, separately for the correct-direction and wrong-direction SNPs. The horizontal thick bars represent the median *F*_*ST*_ of the target SNPs. One data point represents one re-sampling of background SNPs with the same magnitudes of allele frequency as the target SNPs. For both panels, from left to right are traits seed size (SS), ovule number per fruit (ONF), and germination rate (Germ). Asterisks represent the target SNP *F*_*ST*_ is in the upper or lower 5% tail of the background SNPs.

The average genomic divergence between relicts and non-relicts was about 7% of *thaliana-lyrata* divergence [5]. Yet, the mean pairwise differences across haplogroups (Dxy) appear older (Fig. 4c). While haplogroup B-R might be relatively young, it exists in distinct relict groups across Africa and southern Europe (Fig. 4b), whose divergence predated the origin of non-relicts [4, 6]. The results suggested that the divergent evolution of seed sizes did not result from the rapid fixation of novel mutations in either relicts or non-relicts but was likely due to the divergent sorting of standing variations during the allopatric phase. Upon secondary contact, the haplogroups conferring smaller seeds (B-N and C) expanded across Eurasia with non-relicts. Consistently, no obvious trace of extended haplotype homozygosity was identified for analyses comparing relicts versus non-relicts (Fig. 4d). Among haplogroups, only group C had slightly higher extended homozygosity.

### Signatures of polygenic adaptation on single traits

Since most variations in these crucial traits are explained by polygenic factors, we investigated whether traces of divergence selection exist in these polygenes. For each of the ∼240k LD-pruned SNPs, we designated the allele with higher frequency in relicts than in non-relicts as the “relict allele” and the other as the “non-relict allele.”

To investigate the trait effect of SNPs with the highest divergence, GWAS scores were first polarized so that SNPs whose relict allele decreases trait value have negative scores. The GWAS scores of SNPs with top 1% *F*_*ST*_ values are significantly higher than genomic background for SS and lower for ONF and germination (Fig. 5a), demonstrating that highly diverged loci have stronger effects and correct directions on trait divergence.

To investigate the divergence of SNPs with top GWAS scores, SNPs were first separated into two groups based on their effect directions. For example, SNPs whose relict allele increases SS (with positive signed GWAS scores) would have the “correct direction”, and the opposite for the “wrong direction” SNPs. For ONF and germination, SNPs with negative signed GWAS scores have the correct direction. Under divergent selection, the correct-direction SNPs should be highly diverged, and the wrong-direction SNPs should have allele frequency differences similar to or even less than neutral expectation. The median *F*_*ST*_ of the top 0.01%, 0.1%, and 1% GWAS SNPs were compared with randomly re-sampled background SNPs with similar allele frequencies (see Methods). For the correct-direction SNPs, the top 0.01% have higher *F*_*ST*_ than genomic background for SS but not the two other traits (Fig. 5b), suggesting the top SNPs indeed had higher divergence in the Mendelian trait SS. The top 0.1% and 1% SNPs have higher *F*_*ST*_ than the genomic background in all traits, consistent with natural selection driving the allele frequency divergence of many minor-effect loci. For the wrong-direction SNPs, in most cases, the *F*_*ST*_ was either similar to or significantly lower than the genomic background, implying that natural selection did not act on them or even prevented their divergence.

We further utilized a method to estimate whether the among-population quantitative trait divergence is higher than the neutral expectation, using the combined effects of polyloci [24, 25]. Accessions from Eurasia and Africa were separated into 21 populations following previous studies [6]. For all three traits, the Q_*X*_ values computed from the effect sizes and allele frequencies of GWAS top 100 SNPs are significantly larger than those from background SNPs (Supplementary Fig. 6), consistent with our results that divergent selection has driven the quantitative trait differences among populations.

**Figure 6.**
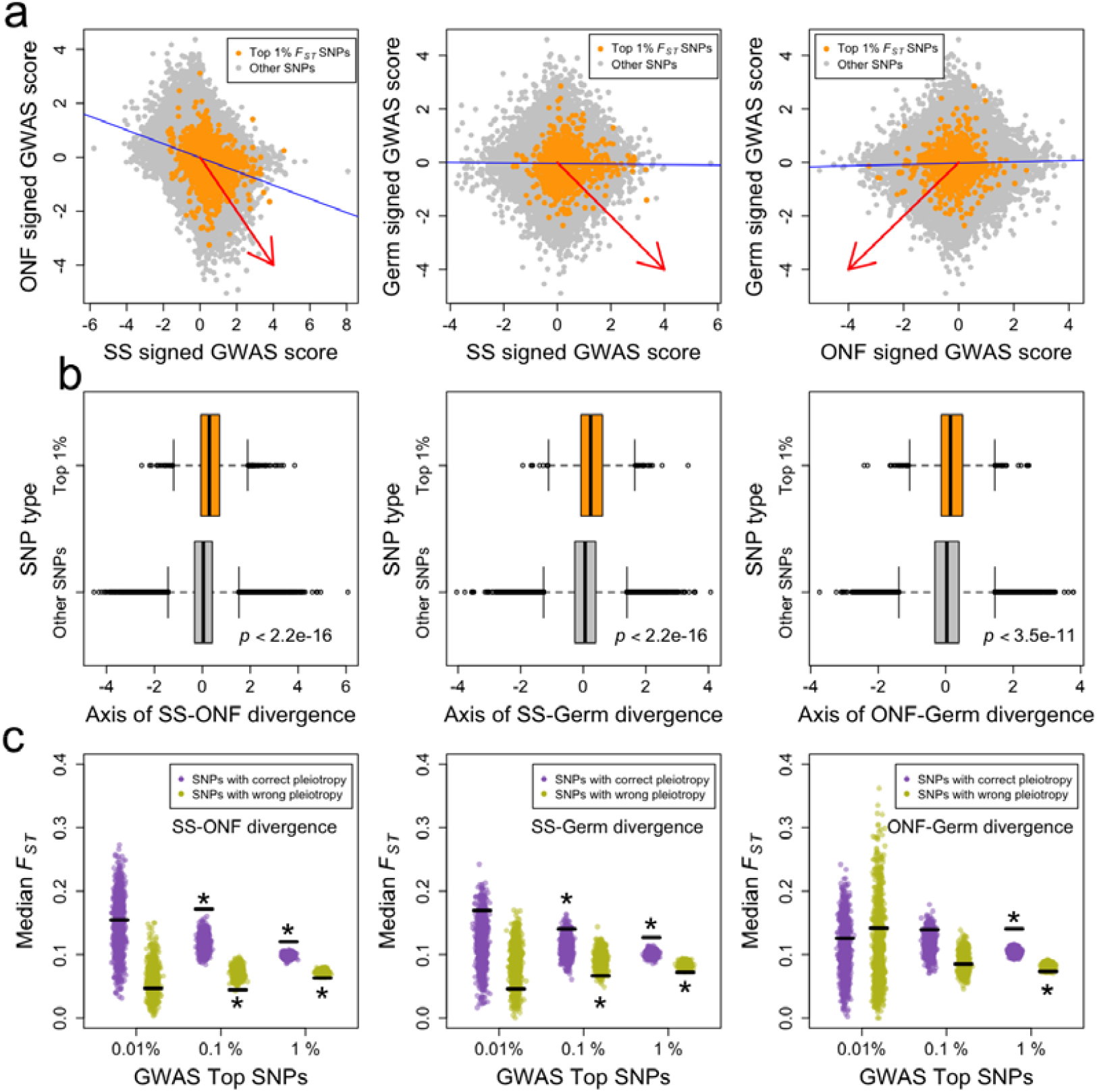
Signatures of polygenic adaptation on trait covariances. **(a)** Scatter plots of SNP GWAS scores on pairs of traits. The GWAS scores were polarized so that SNPs whose relict allele decreases trait values have negative values. The blue lines are regression lines, and the red arrow points to the correct quadrant where SNPs have the correct directions in this multi-trait relationship. The projection coordinates of SNPs on the axis defined by the red arrow are the composite GWAS scores for **(b)** and **(c). (b)** Comparison of composite GWAS scores for SNPs with the top 1% *F*_*ST*_ value versus other SNPs. **(c)** Comparison of *F*_*ST*_ values for SNPs with the top 0.01%, 0.1%, or 1% composite GWAS scores, separately for the correct-direction and wrong-direction SNPs. The horizontal thick bars represent the median *F*_*ST*_ of the target SNPs. One data point represents one re-sampling of background SNPs with the same magnitudes of allele frequency as the target SNPs. Asterisks represent the target SNP *F*_*ST*_ is in the upper or lower 5% tail of the background SNPs.

### Signatures of polygenic adaptation on trait covariances

Given the complex inter-connected regulation networks in a cell and the potentially widespread “network pleiotropy” [26], we investigated whether SNPs with large effects and correct directions simultaneously on multiple traits would be favored by selection. Similar to single-trait analyses, SNP GWAS scores were polarized to reflect the effect direction of the relict allele. Plotting the signed GWAS scores between trait pairs, we found a significant SNP-effect-level correlation in all pairs. SS and ONF have an especially strong negative relationship (Fig. 6a), reflecting the trade-off between seed size and number in a fruit. Similarly, we defined correct-direction SNPs as those in the quadrant with correct directions in both traits (e.g., relict allele increasing SS and decreasing ONF), and SNPs in the three other quadrants have the wrong directions for this multi-trait evolutionary relationship. We further projected all SNPs on the axis pointing towards the correct quadrant (red arrows in Fig. 6a) and used the coordinates as signed composite GWAS scores. For all trait pairs, SNPs with top 1% *F*_*ST*_ values have significantly higher composite GWAS scores than genomic background (Fig. 6b), suggesting highly diverged loci have more substantial effects and correct among-trait directions.

Like the single-trait case, we investigated whether SNPs with the most extreme 0.01%, 0.1%, or 1% composite GWAS scores are highly diverged, separately for the correct- or wrong-direction SNPs. For the top 0.01% SNPs, the median *F*_*ST*_ does not differ from the genomic background, suggesting the patterns of trait covariances were not mediated by Mendelian loci (Fig. 6c). This is consistent with the lack of GWAS peak overlap among these traits (Fig. 2a). In most cases of the top 0.1% or 1% SNPs, the *F*_*ST*_ values are larger for correct-direction and lower for wrong-direction SNPs than background SNPs. The results are consistent with the idea that natural selection did not only drive the divergence of single traits but also the co-evolution of multiple traits through polygenes.

## Discussion

A species is composed of distinct populations, and the composition of a species could be changed by frequent demographic events such as population admixture, subdivision, migration, or extinction. Among them, large demographic turnover is one of the most dramatic, as the exclusion of most populations by a dominant one substantially reduced species-wide genetic variation. While most demographic events could be caused by external factors such as environmental change, demographic turnover likely has a genetic basis, which remains uninvestigated.

Focusing on *Arabidopsis thaliana*, we provide one of the few investigations of the phenotypic and genetic bases of such demographic events. It is commonly believed that weedy plants exhibit greater dispersal capability, prolific seed production, and shorter life cycles [8, 9]. Consistent with this notion, non-relicts produce more and smaller seeds with higher germination rates (Fig. 1), contributing to their higher fecundity, dispersal ability, and colonization potential [27]. Given similar fruit sizes between the two groups, the smaller seeds of non-relict is a trade-off (with a strong polygenic basis, Fig. 6a) to produce more seeds in the same fruit. While other studies suggested that larger seeds might have a higher survival rate than small seeds[28, 29], we did not find that non-relicts perform worse in germination or survival.

Why have relicts not been completely replaced so far? While relicts are restricted to natural, warmer, drier environments[3, 30], the proportion of non-relicts is higher in high-disturbance environments such as artificial and agricultural areas [3]. In addition, some relicts are locally adaptive to specific environments, such as volcanic areas with low soil manganese [31]. It was also shown that pure non-relicts could not completely replace local relicts, as non-relicts inhabiting locations previously occupied by relicts possess introgressions of locally adaptive genomic segments from relicts[5, 6]. Based on the competitor/stress-tolerator/ruderal (CSR) model of plant life history traits [32], non-relicts are more similar to ruderals, while some relicts might be stress-tolerators. In addition to local adaptation, the geographic barrier might be another critical factor as most relicts are currently located in Africa.

The choice of the mapping population is crucial to answering the correct biological question. In this study, we used the Iberian population for GWAS, which is enriched in relict genetic components and is a long-term hybrid zone minimizing problems of population structure [33, 34]. While previous studies also performed GWAS on seed size and ovule number [14, 15], the mapping population consists primarily of non-relicts with little relict genetic variation. As a result, while we identified the Mendelian and polygenic factors contributing to the divergence between relicts and non-relicts, loci identified from previous studies mostly segregate within the non-relict population.

While hard selective sweep has been the focus of many studies, recent opinions emphasized the complex nature of adaptation through quantitative trait evolution [35-37]. We showed the evolution of non-relict weedy traits did not result from novel mutations with large effects. Instead, seed size evolved through the divergent sorting of different ancient haplotypes in a large-effect locus, and the polygenic component of all traits evolved through joint frequency changes in many loci (polygenic adaptation). Interestingly, our novel method demonstrated that polygenic adaptation worked not only in single-trait divergence but also on the relationship between traits with trade-off, with higher allele frequency changes in loci facilitating the adaptive divergence in both traits (Fig. 6). While these complex factors contribute to initial dominance of non-relicts, adaptive introgression is also essential for the successful establishment in novel environments. Therefore, the evolutionary history of weedy *Arabidopsis thaliana* illustrates that different modes of evolution are not mutually exclusive.

This study provides an underlying evolution trajectory resulting in the large demographic turnover, an ecological process we show to have strong genetic bases. The evolution of non-relict weedy traits contributed to their dominance, which could not be simply explained by a few large-effect genes. The results emphasized the importance of previously overlooked factors of ancient genetic variation and polygenic adaptation in driving demographic changes within a species.

## Material and method

### Plant materials

For relict and non-relict comparison, we selected 21 relict and 30 non-relict accessions with little to no introgression from each other[5]. In addition to accessions from Eurasia, the non-relict samples also included a laboratory strain, Col-0, from North America. The relict accessions are mostly from Iberia, Sicily, Lebanon, Cape Verde Islands, and Tanzania.

For the genome-wide association study, we mostly used accessions from the Iberian Peninsula. The Iberian Peninsula is the largest hybrid zone between relicts and non-relicts, and local non-relicts have various levels of introgression from relicts, providing an admixed mapping population to reduce the confounding effect of population structure[5]. Two hundred and fifteen accessions were selected, including 170 Iberian non-relicts, 16 Iberian relicts, 26 non-Iberian non-relicts (including Col-0), and three non-Iberian relicts.

Seeds of all accessions obtained from the stock center were first grown under 22°C and the long day condition (16 hours light: 8 hours darkness) for one generation to bulk up the seeds and eliminate possible maternal effects. The next-generation seeds were used for the following experiments.

### Hypocotyl length measurements

Seeds were sterilized in 70% ethanol with 0.01% Triton X-100 for 10 minutes, followed by a 10-minute wash with 95% ethanol, and then resuspended in 1 ml sterile water. After sterilization, seeds were planted on 0.7% agar gel containing 1x Murashige and Skoog medium (Duchefa, Product code: M0222) with 0.3% sucrose and 0.05% MES (2-(N-Morpholino)ethanesulfonic acid, CAS Number: 145224-94-8) in petri dishes (diameter: 9cm, height: 2cm). The spacing between each seed was at least 0.5 cm to prevent the seedlings from interfering with each other. Each plate contained four accessions and at least 20 seeds of each accession were placed. Plates were kept in the dark at 4°C for one week and then moved to LED growth chambers with the long-day condition (16 hours light: 8 hours darkness) at 22°C. Our experimental procedure follows a previous study investigating the natural genetic variation of hypocotyl response in A. thaliana [38]. At first, the light condition was set to 34 μE/m2/s red light and 7 μE/m2/s blue light to induce germination. After 24 hours, the far-red light was adjusted to let the ratio of red light and far-red light (R: FR) be 2. After 48 hours, the R: FR ratio in shade simulation experiments was lowered to 0.5, while the R: FR ratio in sun simulation experiments was kept at 2 for four days. A spectrometer (HiPoint, HR350) was used to measure the light condition inside the LED growth chamber to adjust the settings to the needed conditions. The entire experiment was replicated twice.

A scanner (MICROTEK, ArtixScan F2) was used to scan all seedlings with a ruler to image files with a resolution of 2000 pixels per inch. The hypocotyl length of each seedling was measured and transformed from pixel to millimeter using ImageJ [39]. The shade avoidance response of each accession was calculated as the hypocotyl length differences between R: FR = 0.5 and R: FR = 2.

### Measurements of fruit and seed-related traits

The phenotyping experiments followed a randomized complete block design with four replications, and each replication contains one individual plant from each accession. Un-sterilized seeds of each accession were suspended in 1 ml sterile water and kept in the dark at 4°C to break dormancy and synchronize germination. After one week, around five seeds of each accession were sown in pots (210 ml) containing sterile soil (peat soil: vermiculite: perlite = 4:3:1) and moved to a growth chamber with long daylight (16 hours light: 8 hours darkness) at 22°C. Twenty pots were placed in a rectangular plastic basin (length x width x height = 45 x 35 x 11 cm) and watered twice a week. When watering, 700 ml H2O was poured into the plastic basin. After an additional week, extra seedlings were removed, so there was only one seedling per pot. The accessions that did not flower at four weeks were moved into a 4°C refrigerator with fluorescent lamps for seven weeks for cold vernalization, while others were kept in the growth room.

Each plant’s total number of fruits (FN) was recorded, and undeveloped fruits were excluded from the measurement. Following the previous study [10], from each plant, we harvested three fruits from the primary inflorescence at positions 6 to 10 from the lowest fruit. The following traits were measured: seed number per fruit (SNF), number of undeveloped ovules per fruit (UOF), ovules number per fruit (ONF), developed percentage (DP), and fruit length (FL). SNF and UOF were counted under a stereomicroscope. ONF was obtained as SNF + UOF, and DP is (SNF / ONF) × 100%. Each plant’s seed number per millimeter (SNM) and ovule number per millimeter (ONM) were estimated respectively as SNF/FL and ONF/FL. Each plant’s total seed number (SN) and ovule number (ON) were calculated as SNF × FN and ONF × FN, respectively. For seed size measurement, around 20 seeds from each plant were scanned with the positive film scanner (MICROTEK, ARTIXSCAN F2) [40]. The resolution of each image is 2000 pixels per inch. All image files of the seeds were analyzed using the particle analysis function of ImageJ, and the following measurements were recorded: area (seed size, SS), maximum Feret diameter (seed length, SL), and minimum Feret diameter (seed width, SW). Because these traits were measured from image files, we used pixel as the seed size and length unit.

### *F*_*ST*_ – *Q*_*ST*_ comparisons and heritability

The same set of 51 accessions used in relict and non-relict comparisons were also used for *F*_*ST*_ – *Q*_*ST*_ comparisons. For *F*_*ST*_ estimation, Single-nucleotide polymorphism (SNP) data from the 1001 genome project was used as genotype data. The “--weir-fst-pop” options in VCFtools were used to calculate the *F*_*ST*_ of each SNP between relicts and non-relicts.

*Q*_*ST*_ measures the amount of heritable trait variance among populations relative to the total trait variance [41], similar to *F*_*ST*_. For *Q*_*ST*_ estimation, we used genetic groups (relict and non-relict) and accessions nested within groups as random effects to estimate their variance components with JMP 13 (SAS, Cary, NC, USA). *Q*_*ST*_ of each trait was calculated as

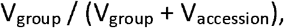

where V_group_ and V_accession_ represent the variance components of the genetic population and accession term, respectively. In outcrossing species, the accession variance component is usually multiplied by two in a typical *Q*_*ST*_ formula. Since A. thaliana is a highly self-fertilizing species and all accessions have been self-fertilized in the stock centers and labs for many generations, it can be modeled as haploid in our calculations [42], as has been done in many A. thaliana studies [43, 44].

The heritability of each trait was calculated as

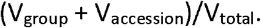

### Genome-wide association study

The least squares mean of each trait for each accession was calculated with JMP 13 (SAS, Cary, NC, USA). The SNP genotype data was obtained from the 1001 genomes project [3].

GEMMA [45] was used for genome-wide association studies. VCFtools was used for biallelic SNPs and individual filtering of genotypic data. PLINK software[46] was used to convert the variant call format (VCF) into PLINK binary ped format, which GEMMA can recognize. The LD-pruned SNP dataset was also generated from PLINK with the “--indep 50 5 2” parameter. The GWAS was conducted using a univariate mixed linear model method with the parameter “-lmm 4”. The relatedness matrix used to control population structure was generated from GEMMA with the “-gk 1” parameter, which calculates the centered relatedness matrix.

The ridge and lasso regressions were calculated using the R package glmnet [47]. Since missing data were not allowed in the analysis, the SNP data were converted to numeric format and imputed by TASSEL 5. The optimal hyperparameter involved in ridge and lasso regression was obtained by fivefold cross-validation. The average cumulative effect (r^2^) between measured and predicted values was acquired by 100 times repeats. Using GWAS results of the 215 accessions, we employed the BSLMM model in GEMMA to predict trait values of 1135 genomes.

### Genetic variance component estimation

We performed a Bayesian Sparse Linear Mixed Model (BSLMM) in GEMMA to estimate the genetic variance components of the traits [48]. The BSLMM algorithm estimates overall heritability as the proportion of phenotypic variance explained (PVE) by the marker and kinship effects. In addition, PGE, the proportion of marker variance (PVE) explained by the genetic effect, was also estimated. In brief, PVE indicates the heritability, PVE x PGE estimates the proportional phenotypic variances explained by the fixed SNP effect, and PVE x (1-PGE) represents the proportional phenotypic variances explained by the random kinship effect. We used 2,000,000 burn-ins and 1,000,000 sampling steps and recorded one state every 10 steps. The median of the 100,000 records represented the results of the estimations. Moreover, we also performed the BSLMM for each chromosome arm following the same procedures. A linear regression was performed using heritability as the response and SNP numbers on each arm as an independent variable.

### DNA isolation

Total genomic DNA for PCR was extracted using a simplified protocol. 100ml extraction buffer used in this protocol was prepared with 1M Tris-HCl 20 ml, 5M NaCl 5 ml, 20% SDS 2.5 ml, 0.5M EDTA 5 ml, and ddH_2_O 67.5 ml. Mature leaves were collected in a microcentrifuge tube with 400μl extraction buffer and ground with micropestel. Plant residues were precipitated by centrifugation at 14000 RPM for 10 minutes. The supernatant was transferred to a new microcentrifuge tube with a 1:1 volume of isopropanol and mixed immediately. The pellet with DNA was harvested by centrifugation at 13000 RPM for 3 minutes, then discarded the supernatant and washed with 75% ethanol. After the supernatant was discarded, we placed the tubes in a fume hood until the ethanol evaporated. DNA samples were dissolved in nuclease-free water for further experiments.

### T-DNA insertion lines screening

T-DNA insertion lines of the genes

(*AT2G32520, AT2G32530, AT2G32540, AT2G32560, AT2G32580, AT2G32590, AT2G32 600*) under our seed size GWAS peaks were ordered from Arabidopsis biological resource center (ABRC). Around 10 seeds of each mutant line were planted in pots to bulk up and confirm the genotype of each individual. We used PCR for T-DNA verification, and three primers were designed: genomic primers on either side of the T-DNA insertion point and a primer on the T-DNA. Wild-type individuals will only get the PCR product with the genomic primer pairs. Homozygous mutant lines will only get the product with the primer on the T-DNA insertion and the reverse genomic primer. For heterozygous lines, we can get PCR products from both primer pairs. Seeds were collected from the homozygous mutant individual for further experiments. Since T-DNA insertion lines of *AT2G32590* can’t get homozygous mutant individuals (likely mutant homozygous lethal), only seeds from heterozygous individuals were collected.

### RNA isolation and quantitative real-time qPCR analysis

Seeds collected from heterozygous SALK-017766 and SALK-069185 lines were planted for RNA extraction from buds and young siliques. Before sample collection, the genotype of each individual was checked using general PCR. Tubes with zirconia/silica beads were used for sample collection. Buds or young siliques from four individuals with the same genotype were pooled in the same tube as a biological replicate. Each tissue and genotype had four biological replicates for qPCR. Plant tissues were disrupted by homogenizer, and total RNA was extracted using the QIAGEN RNeasy plant mini kit (QIAGEN, Inc., Valencia, CA). We removed the genomic DNA from RNA samples by DNase I (New England Biolabs, Cat. M0303). After the treatment of DNase I, a 1:1 volume of 8M LiCl was added, and the samples were stored at -20 °C for at least 2 hours. The pellet with RNA was harvested by centrifugation at 13000 RPM for 40 minutes at 4°C and washed with cold 80% and 100% ethanol. We discarded the supernatant and dried the pellet in the fume hood for about 5 minutes. RNA samples were dissolved in nuclease-free water. RNA concentration and the A260/A280 A260/230 ratio were measured by NanoDropTM One. SuperScript™ III Reverse Transcriptase (Thermo Fisher Scientific, Waltham, MA, USA) was used for first-strand cDNA synthesis from a fixed concentration of RNA samples. RT-qPCR was performed with CFX Connect or CFX384 Touch Real-Time PCR Detection System (Bio-Rad) using iQ™ SYBR^®^ Green Supermix.

### GUS staining

X-Gluc 10 mg was dissolved in 1 mL methanol to prepare X-Gluc stock. The X-gluc substrate solution was made with 100 μl X-Gluc stock, 1 mL 2x phosphate buffer pH 7.0 (made of 0.1 M NaH_2_PO_4_ and 0.1 M Na_2_HPO_4_), 20 μL 0.1 M potassium ferrocyanide, 20 μL 0.1 M potassium ferricyanide, 10 μL 10 % Triton X-100 and 0.85 mL water. The plant materials to be tested were first fixed in 90% acetone for 20 minutes on ice and then rinsed with 1x phosphate buffer[49, 50]. The plant tissues were stained in X-gluc substrate solution at 37°C overnight. After staining, we used the improved Hoyer’s method [51] for removing the chlorophyll and tissue clearing. The specimens were placed on slides after clearing and examined under a microscope.

### The cell size of the embryo

Mature dried seeds were soaked in H_2_O for 1 hour and then dissected under the stereomicroscope to isolate mature embryos from seed coats [52]. The isolated embryos were fixed with ethanol:acetic acid (3:1) for at least 1 hr. After fixation, EtAc in the tube was replaced with the improved Hoyer’s medium (chloral hydrate: distilled water: glycerol = 10:6:1 g) for tissue clearing for 1 hour [51, 53]. Observation and photos of the embryo were taken using an Olympus compound microscope with a differential interference contrast (DIC) option. The area of the cotyledon, palisade and spongy cells were measured by ImageJ.

### Test of polygenic adaptation

To test the polygenic adaptation in single traits, we compared the *F*_*ST*_ of candidate SNPs versus the genomic background using the LD-pruned SNP dataset with around 240k SNPs. The *F*_*ST*_ of each SNP was calculated using 99 non-relicts and 100 relicts. For each bi-allelic SNP, we first designated an allele to be the “relict allele” if its allele frequency is higher in the relicts than the non-relicts. SNP GWAS scores were then polarized to reflect the effect direction of the relict allele, and SNPs whose relict allele decreases trait value would have negative GWAS scores. The signed GWAS scores were compared between SNPs with top 1% *F*_*ST*_ values and other SNPs to investigate whether highly diverged SNPs tend to have stronger effects and correct directions on traits.

To investigate whether SNPs with strong effects are highly diverged, SNPs were first separated into two groups (with correct or wrong directions) based on the effect directions. A SNP was categorized as the “correct-direction SNP” if the relict allele affects the trait in a direction consistent with population divergence (e.g., the relict allele increases seed size, decreases ovule number per fruit, or decreases germination rate), and a “wrong-direction SNP” has the opposite effect. The *F*_*ST*_ comparisons were performed separately for these two types of SNPs. Following previous studies [54, 55], we compared the levels of divergence between target SNPs versus background SNPs (both set already LD-pruned) while controlling for their minor allele frequency (MAF). Specifically, all SNPs were first separated into ten categories based on their MAF, with a 0.05 bin size. The target SNPs with the 0.01%, 0.1%, or 1% highest GWAS scores were also separated into these MAF bins. For each re-sampling, genomic SNPs with amounts equal to the target SNP were sampled from the same MAF bins, and their median *F*_*ST*_ value was recorded. The re-sampling was repeated 1,000 times to obtain a background distribution of median *F*_*ST*_, representing the levels of divergence if one randomly samples the same number of SNPs with similar MAF distribution as the target SNPs. The median *F*_*ST*_ of the top GWAS SNPs was then compared with this background distribution.

We further conducted the *Q*_X_ test to estimate levels of among-population trait divergence [24, 25]. Instead of comparing SNP *F*_*ST*_, this method uses SNP effect sizes and population allele frequencies to calculate the expected population trait value. Similar to our approach above, this method first uses SNPs with top GWAS scores to estimate the population trait value and compares it with those calculated from genomic background SNPs with similar minor allele frequencies and LD patterns. This analysis used the same LD-pruned SNP dataset and GWAS results as in the SNP *F*_*ST*_ comparison. The accessions used in *Q*_*X*_ test were separated into different regions and clusters based on their geographical location and genetic structure. We followed the methodology outlined in the article by Hsu, Che-Wei *et al*. [6] to group the accessions in Europe. Relicts from Morocco were categorized into groups according to the study conducted by Durvasula, Arun *et al*. [4]. Allele frequency was calculated using the “--ferq” option in VCFtools.

To investigate the polygenic adaptation on trait covariances, we used the same dataset to perform similar permutation-based analyses. The SNPs were first polarized based on the effect direction of the “relict allele” on a trait, and the GWAS score (-log_10_ *P* value) of SNPs whose relict alleles decrease trait values would be designated as negative. Plotting the signed GWAS scores of two traits, one could investigate the SNP-level genetic correlation between the two traits. In this two-dimensional space of the signed GWAS score of two traits (where each data point is a SNP), we defined the “axis of population divergence” as the axis pointing towards the quadrant where SNPs have the correct direction on both traits (e.g., relict alleles increasing SS and decreasing ONF). All SNPs were projected onto this axis, and their coordinates on the axis are the new “signed composite GWAS scores” of this trait combination.

Along this axis of population divergence, we compared whether SNPs with the 1% highest *F*_*ST*_ values have higher signed composite GWAS scores than the rest. Based on this between-trait relationship, a SNP could be categorized as the “correct-direction SNP” if it exists in the quadrant with the correct directions of divergence (e.g., the relict allele has a positive signed GWAS score on seed size and negative signed GWAS score on ovule number per fruit). Otherwise, it would be categorized as the “wrong-direction SNP.” This is the same logic we used previously, except being applied to the association between two traits. Like the single-trait procedure, the permutation-based method was used to investigate whether SNPs with the top 0.01%, 0.1%, or 1% composite GWAS scores have higher median *F*_*ST*_ values than background SNPs with similar minor allele frequency distributions. In this case, the top wrong-direction SNPs are those with the most negative signed composite GWAS scores.

## Acknowledgment

We thank Che-Wei Hsu, Yang-Tui Cheng, and Yi-Fang Tsay’s lab for their assistance and Magnus Nordborg and Tal Dahan for comments on the manuscript. We are grateful to the support from National Taiwan University’s Computer and Information Networking Center for high-performance computing facilities as well as Technology Commons of College of Life Science for molecular biology assistance. C.-R.L. was funded by 105-2311-B-002-040-MY2 and 111-2628-B-002-021 from the Ministry of Science and Technology, Taiwan.

